## Supplementary Information for "GRN_modeler: An Intuitive Tool for Constructing and Evaluating Gene Regulatory Networks and its Applications to Oscillators and a Light Biosensor"

#### 1 Experiments

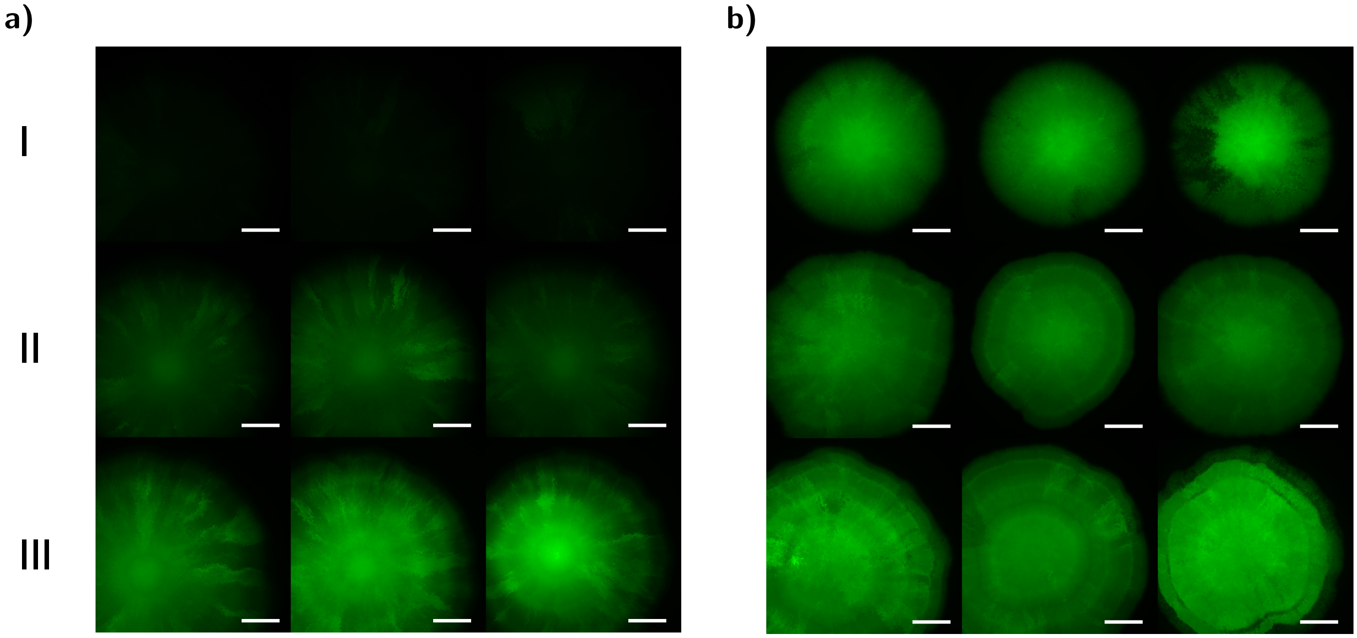


**Figure S1:** Fluorescence microscopy images of colonies harboring the light-inducible 1-node circuit (pJP_1node and pJP_Bla01). Colonies grew for 4 days under square wave light pulses (T = 24h) and 50% duty cycle. a - Cultures without L-arabinose. b - Cultures with 0.2% L-arabinose. I. No light exposure, II - light pulses with 25% of the maximal intensity of the LITOS device, III - light pulses with 100% of the maximal intensity of the LITOS device. Scale bars = 1 mm.

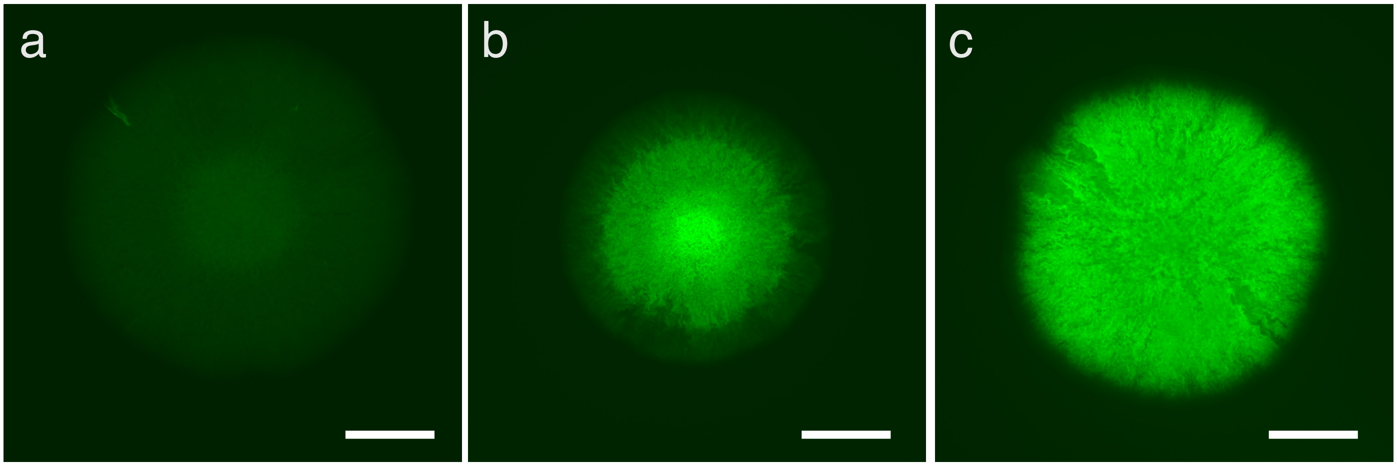


**Figure S2:** The light-inducible CRISPRlator (pJP_Osc05, pJP_Bla01 and pJ1996_v2) is not oscillating on solid surface. Fluorescence microscopy pictures of colonies grown for 5 days in dark condition (a), at constant light of 40% intensity (b) and in the dark with 0.2% L-arabinose (c). Scale bars = 1 mm.

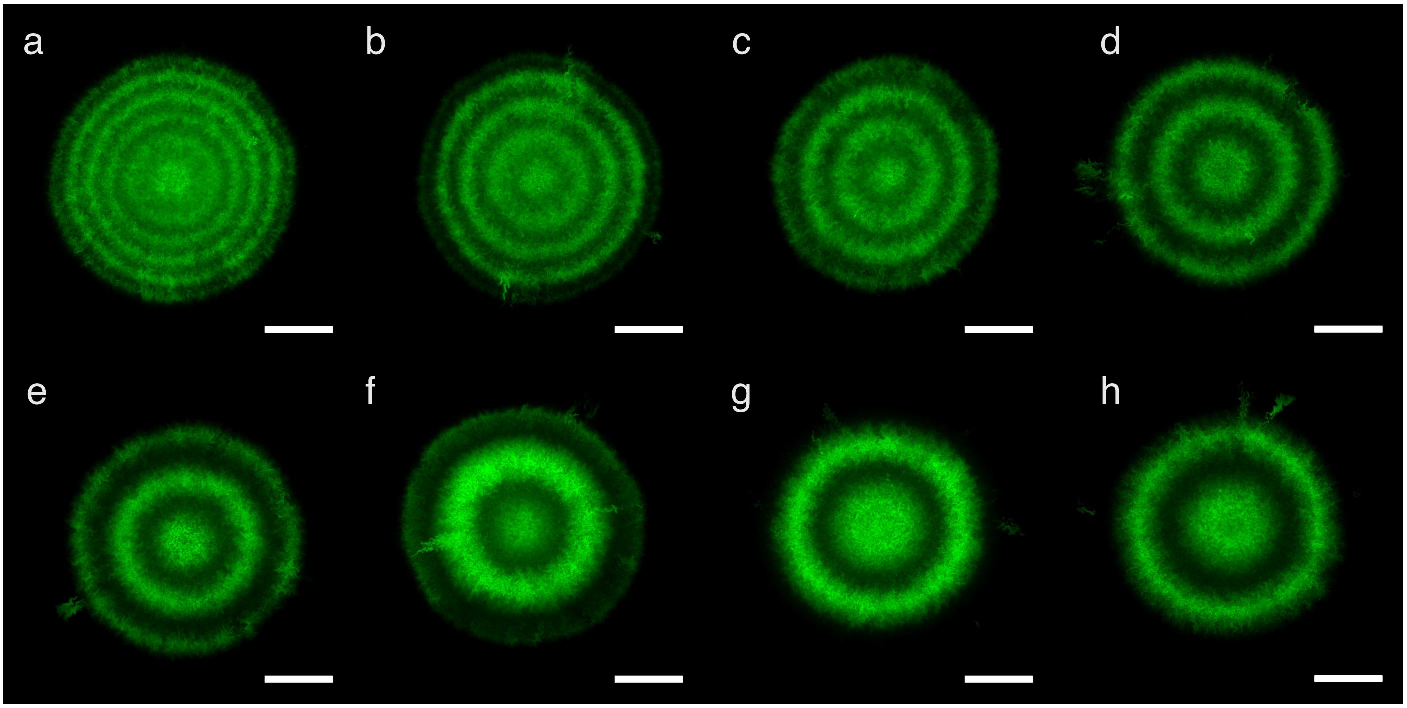


**Figure S3:** The number of rings generated by the light-inducible CRISPRlator matches with the number of light pulses. Fluorescence microscopy pictures of colonies harboring the light-inducible CRISPRlator (pJP_Osc05, pJP_Bla01 and pJ1996_v2). mCitrine is represented in green. The colonies grew for 4 days under square wave light pulses (a - 12h, b - 14h, c - 16h, d - 18h, e - 20h, f - 22, g - 24h, h - 26h), with a duty-cycle of 50%. Scale bars = 1 mm.

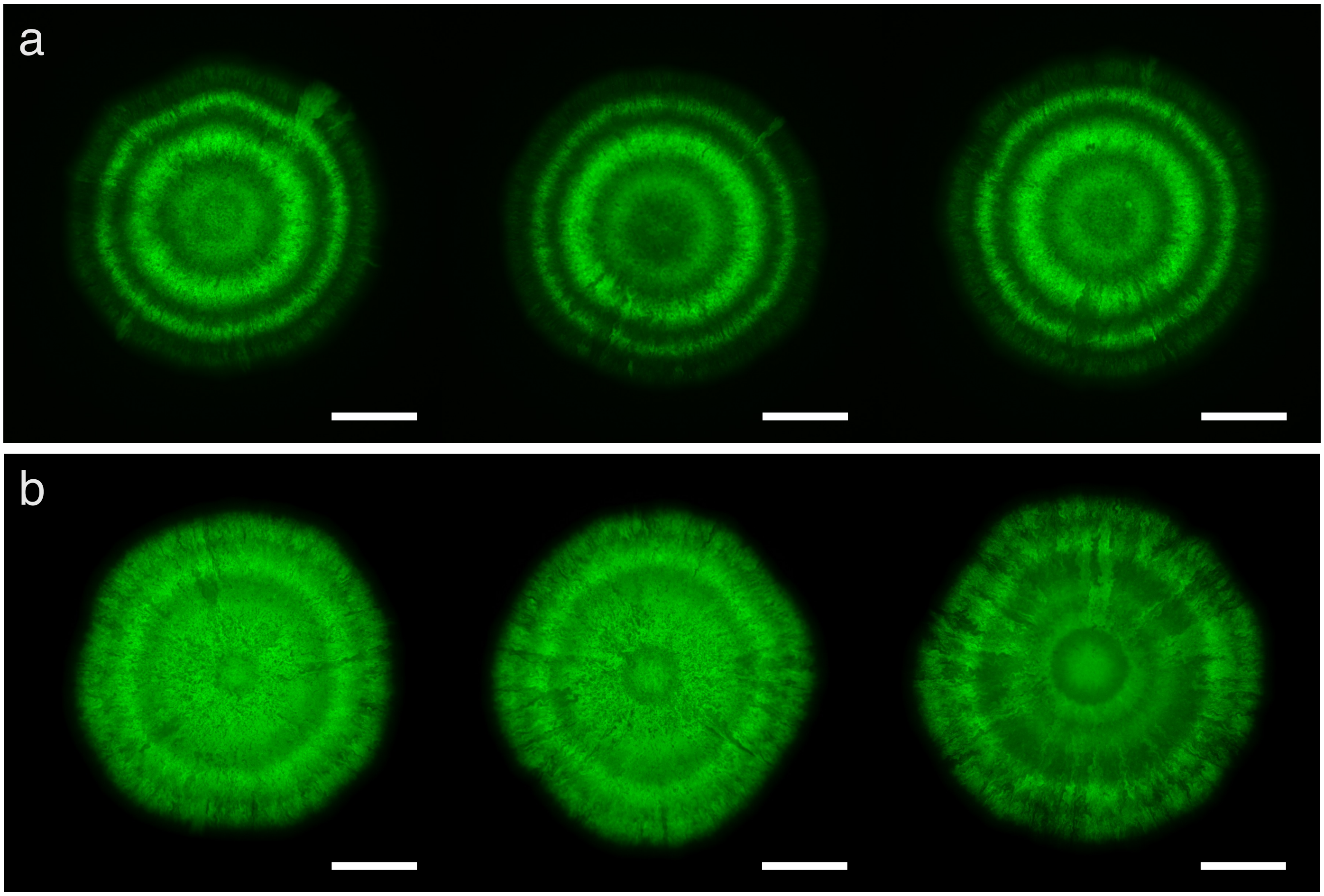


**Figure S4:** Phase and anti-phase ring patterns by the light-inducible CRISPRlator (pJP_Osc05, pJP_Bla01 and pJ1996_v2). a) Three replicates of colonies grown on solid surface and subjected to light pulses from 0% to 15% intensity (41.66% duty-cycle) for 4 days. b) Colonies grown under the same condition as in (a), but with addition of 0.2% L-arabinose. Scale bars = 1 mm.

| Plasmid name | Description | Reference Number (Addgene) | Reference |
|---|---|---|---|
| pBLADE_ ONLY_C | Plasmid encoding the light-inducible VVD-AraC. | #168050 | [1] |
| pJP_Bla01 | Modified pBLADE_ ONLY_C, with gentamicin resistance gene and deleted f1 origin of replication. | #230982 | This work |
| pJ1996_v2 | Contains *dCas9* and *csy4* necessary for CRISPRi. | #140664 | [2] |
| pJ2072 (1-OS2) | L-arabinose inducible CRISPRlator, it is composed of three nodes, with each node producing a sgRNA and a fluorescent reporter. It was used as backbone to construct pJP_Osc05. | – | [3] |
| pJP_Osc05 | pJ2072 (1-OS2) without *araC* and Kanamycin resistance gene placed between mCitrine and mCerulean nodes. | #229040 | This work |
| pCOLA-AraC-pBad-GFP | colA backbone with pBad promoter controlling the expression of sfGFP. | | [4] |
| pJP_Ctrl04 | pCOLA-AraC-pBad-GFP without *araC*. | #230983 | This work |
| pJP_1Node | pJP_Ctrl04 with pBad controlling the expression of mCitrine instead of sfGFP. | #230984 | This work |

**Table S1:** List of plasmids used in this work.

### 2 Simulations

#### 2.1 Command line functionality

- `Cell('ModelName')`: Instantiates a new object for the gene regulatory network specified by `'ModelName'`. This name should match one of the models implemented in the **models** folder. The currently available models are "Elowitz," "Tomazou," and "CRISPR," as described in the Results section. For example, use `Ecoli = ... Cell('Elowitz');` to create a new instance of the "Elowitz" model with the user-defined name "Ecoli".

- `add_node('NodeName', 'NodeType')`: This method adds a new node to the system with the specified `'NodeName'` and `'NodeType'`. If only one node type is defined in the model, specifying the type is optional, and a simplified naming convention is used for parameters that do not include the type. This simplification also applies to proteases and regulators. For example, use `Ecoli = Ecoli.add_node('N1', 'type1');` to add a node named 'N1' with type 'type1' to the `Ecoli` model.

- `add_protease('NodeName', 'ProteaseName')`: Adds a protease, specified by `'ProteaseName'`, to the node identified as `'NodeName'`. For example, use `Ecoli = Ecoli.add_protease('N1', 'PROT1', 'type1');` to add the protease `'PROT1'` to the node `'N1'` in the `Ecoli` model. The purpose of this function is to introduce an additional species that can interact with existing species across different nodes. We demonstrate this feature using a protease as an example, but it could also be applied to RNA or other species.

- `add_regulator('RegulationType', 'obj_name', 'obj_input', 'reg_name1')`: Adds a regulator to a specified object. In Figure S5 we show the general concept of the regulation. The regulation type is defined by `'RegulationType'`. The name of the regulated object is `'obj_name'`, and its selected input is `'obj_input'`, which is regulated by `'reg_name1'`. For example, to add a `'Repression_out'` type regulation to the `'N1'` node through its input named `'HILL'`, regulated by the `'R1'` species (`N1HILL|-R1`), use: `Ecoli = ... Ecoli.add_regulator('Repression_out', 'N1', 'HILL', 'R1');`. To add regulators to existing regulations, we can extend the input list using the following syntax: `add_regulator('RegulationType', 'obj_name', ... 'obj_input', 'reg_name1', 'reg_input1', 'reg_name2')`. Here, `'reg_input1'` specifies the input of the first regulator (`'reg_name1'`), while `'reg_name2'` is the name of the new regulator. For example, to add another regulator (`'R2'`) to the existing regulation (`N1HILL|-R1`) with a type `'Activation_out'` for the `'HILL'` input (`N1HILL|-R1HILL<-R2`), use: `Ecoli = Ecoli.add_regulator('Activation_out', 'N1', 'HILL', 'R1', ... 'HILL', 'R2');`. If only one input function is implemented in the model, this can be simplified to: `Ecoli = ... Ecoli.add_regulator('Activation_out', 'N1', 'R1', 'R2');`. If there is only one input, the naming process will be simplified both in the graph and in the variable names, resulting in a representation like `N1|-R1<-R2`. Regulators can have an unlimited number of hierarchical levels by further extending the input list in the same way. To simplify this process, one can utilize the code generation feature of the GUI.

- `set('Parameter_name', 'Property', Value, 'object_name')`: Sets the property (`'Property'`) of a specified parameter (`'Parameter_name'`) to a given value (`Value`) within a specific object (`'object_name'`). If the parameter belongs to a regulator, you can extend the input list with additional regulator names as needed. For example, `Ecoli.set('P_N2', 'InitialAmount', 100, 'N2');` sets the initial amount of the species `'P_N2'` to `100` for the `'N2'` node.

- `get('Parameter_name', 'Property', 'object_name')`: This method retrieves the property (`'Property'`) of a specified parameter (`'Parameter_name'`) for a given object (`'object_name'`). The syntax is similar to the `set` method, but the `Value` is the output rather than the property being set. For example, `Value = ... Ecoli.get('P_N2', 'InitialAmount', 'N2');` retrieves the initial amount of the parameter `'P_N2'` for the object `'N2'`.

- `make_graph()`: Generates a graph for the GRN, where the nodes, proteases, and regulators are the "dots" of the graph, and the interactions are represented by arrows in the directed graph. For example, use `Ecoli.make_graph();` to create the graph for the `Ecoli` model.

- `get_model()`: Creates a SimBiology model object. This model can be opened with the SimBiology application for analysis or to run simulations. For example, use `Mobj = Ecoli.get_model();` to create a model object from the `Ecoli` instance.

- `[t, c, names] = run_simulation(model, solver)`: This function runs a simulation on the specified `model` using the `solver`. If the `solver` is a COPASI solver rather than a built-in MATLAB solver, this command can be used similarly to MATLAB's `sbiosimulate` function. Here, `t` represents the output time points, `c` is a matrix of concentrations, and `names` contains the species names. The simulation length, tolerance values, and tracked species can be configured using SimBiology's built-in functionalities, such as the `getconfigset` function. To run a stochastic simulation, the reactions should be rewritten to create a COPASI-compatible SBML file:

```
configset = getconfigset(Mobj); %Retrieve the simulation settings from the model
```

```
Mobj = convert2irrev(Mobj); %Convert reversible reactions into two irreversible reactions
Mobj = correct_modifiers(Mobj); %Rewrite the reactions to make them COPASI compatible
Ecoli.set_configset(Mobj, configset); %Apply the original configuration settings
[t, c, names] = run_simulation(Mobj, 'adaptivesa'); %Run the simulation with the selected ...
adaptivesa solver
```

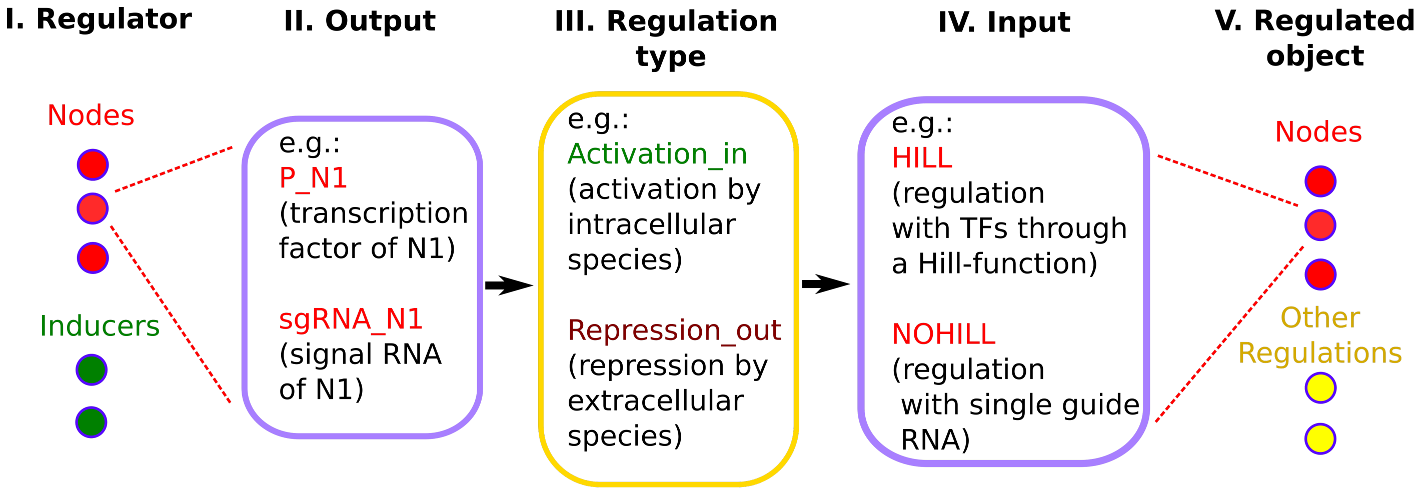


**Figure S5:** The basic concept and structure for setting up the regulatory system. Regulators (such as nodes or inducers like L-arabinose) can have multiple outputs (e.g., transcription factors or sgRNA) to control different targets. After selecting the appropriate regulation type, the regulator can be connected to a regulated object (another node or regulation) via its input, forming the regulatory network.

#### 2.2 Elowitz-model

The specifics of the Elowitz model are provided in Table S2. In reaction $R_1$, the $i$th node generates mRNA, which can be inhibited by a transcription factor originating from the $j$th node. Reaction $R_2$ describes the production of the transcription factor associated with the $i$th node.

**Table S2:** The Elowitz-type transcription factor model [5] describes the production of mRNA and protein ($\mathrm{mRNA}_i$, $\mathrm{P}_i$) at the $i$th node, which is repressed by another transcription factor, $\mathrm{P}_j$. This repression is modeled using a Hill function: $\mathrm{HILL}([\mathrm{P}_j]) = \frac{1}{1+([\mathrm{P}_j]/\mathrm{K})^n}$, where $\mathrm{K} = 40$ molecule and $\mathrm{n} = 2$.

| Nr. | Reaction | Rate law | Rate constant [5] | Unit |
|---|---|---|---|---|
| $R_1$ | $\emptyset \longleftrightarrow \mathrm{mRNA}_i$ | $r_1 = k_0 + k_1 \cdot \mathrm{HILL}([\mathrm{P}_j])$ | $k_0 = 0.03$ | molecule/minute |
| | | | $k_1 = 30$ | molecule/minute |
| | | $r_{1r} = k_2\,[\mathrm{mRNA}_i]$ | $k_2 = 0.3466$ | 1/ minute |
| $R_2$ | $\emptyset \longleftrightarrow \mathrm{P}_i$ | $r_2 = k_3\,[\mathrm{mRNA}_i]$ | $k_3 = 6.9315$ | 1/ minute |
| | | $r_{2r} = k_4\,[\mathrm{P}_i]$ | $k_4 = 0.0693$ | 1/ minute |

Here we showcase how to create a model for the repressilator with our tool using the command line functionalities:

**Listing 1:** Building the model for the Repressilator

```
Ecoli = Cell('Elowitz');
Ecoli = Ecoli.add_node('N1','type1');
Ecoli = Ecoli.add_node('N2','type1');
Ecoli = Ecoli.add_node('N3','type1');
Ecoli = Ecoli.add_regulator('Repression_in','N2','HILL','P_N1');
Ecoli = Ecoli.add_regulator('Repression_in','N3','HILL','P_N2');
Ecoli = Ecoli.add_regulator('Repression_in','N1','HILL','P_N3');
```

#### 2.3 Tomazou-model

In Table S3 we present the Tomazou-type transcription factor model [6]. The mRNA production of the $i$th node can be repressed by the transcription factor of the $j$th node, $\mathrm{P}_j$. This repression is described with the following Hill-function: $\mathrm{HILL}([\mathrm{P}_j]) = \frac{1}{1+([\mathrm{P}_j]/\mathrm{K})^n}$, where K = 5 molecule and n = 2. When external inducers are present ($\mathrm{I}_1$ and $\mathrm{I}_2$ in Figure 3II-a,b in the manuscript, $[\mathrm{P}_j]$ can be replaced with the concentration of the inducers in the Hill-function and K = 50 $\mu$M. The protease degradation rate is determined using Michaelis-Menten kinetics: $\mathrm{k}_{\mathrm{protease}} = \frac{\mathrm{k}_{\mathrm{protease,max}}[\mathrm{PROT}]}{\mathrm{K}_{\mathrm{protease}} + \mathrm{Substrates}}$, where $\mathrm{k}_{\mathrm{protease,max}} = 50$ 1/ minute, $\mathrm{K}_{\mathrm{protease}} = 30$ molecule and "Substrates" represents the sum of all proteins degraded by the given protease ($\mathrm{Substrates} = \sum_i [\mathrm{P}_i]$).

**Table S3:** The Tomazou-type transcription factor model [6]. Further explanation can be found in the text.

| Nr. | Reaction | Rate law | Rate constant [6] | Unit |
|---|---|---|---|---|
| $\mathrm{R}_1$ | $\emptyset \longleftrightarrow \mathrm{mRNA}_i$ | $\mathrm{r}_1 = \mathrm{n}_{\mathrm{copy}}(\mathrm{a}_0 + \mathrm{a}_1 \cdot \mathrm{HILL}([\mathrm{P}_j]))$ | $\mathrm{n}_{\mathrm{copy}} = 25$ | molecule |
| | | | $\mathrm{a}_0 = 0.001$ | 1/ minute |
| | | | $\mathrm{a}_1 = 100$ | 1/ minute |
| | | $\mathrm{r}_{1r} = (\mathrm{k}_{\mathrm{mRNA,degr}} + \mathrm{k}_d)\,[\mathrm{mRNA}_i]$ | $\mathrm{k}_{\mathrm{mRNA,degr}} = 0.5$ | 1/ minute |
| $\mathrm{R}_2$ | $\emptyset \longleftrightarrow \mathrm{uP}_i$ | $\mathrm{r}_2 = \mathrm{k}_{\mathrm{translation}}\,[\mathrm{mRNA}_i]$ | $\mathrm{k}_{\mathrm{translation}} = 6.9315$ | 1/ minute |
| | | $\mathrm{r}_{2r} = (\mathrm{k}_d + \mathrm{k}_{\mathrm{protease,i}})\,[\mathrm{uP}_i]$ | $\mathrm{k}_d = 0.01$ | 1/minute |
| $\mathrm{R}_3$ | $\mathrm{uP}_i \longrightarrow \mathrm{P}_i$ | $\mathrm{r}_3 = \mathrm{k}_{\mathrm{mat}}\,[\mathrm{uP}_i]$ | $\mathrm{k}_{\mathrm{mat}} = 0.4$ | 1/ minute |
| $\mathrm{R}_4$ | $\mathrm{P}_i \longrightarrow \emptyset$ | $\mathrm{r}_4 = (\mathrm{k}_d + \mathrm{k}_{\mathrm{protease,i}})\,[\mathrm{P}_i]$ | | |

Here we showcase how to create the model presented in Figure 3II of the manuscript:

**Listing 2:** Building the model for independent amplitude and frequency modulation in a re-designed Repressilator

```
%% Add nodes
Ecoli = Ecoli.add_node('R1');
Ecoli = Ecoli.add_node('R2');
Ecoli = Ecoli.add_node('R3');
Ecoli = Ecoli.add_node('G');

```

```
%% Add protease
Ecoli = Ecoli.add_protease('R1','C');
Ecoli = Ecoli.add_protease('R2','C');
Ecoli = Ecoli.add_protease('R3','C');
Ecoli = Ecoli.add_protease('G','L');

%% Add regulators
Ecoli = Ecoli.add_regulator('Repression_in','R1','R3');
Ecoli = Ecoli.add_regulator('Repression_in','R2','R1');
Ecoli = Ecoli.add_regulator('Repression_in','R3','R2');
Ecoli = Ecoli.add_regulator('Repression_in','G','R3');
Ecoli = Ecoli.add_regulator('Activation_in','R2','Y');
Ecoli.set('Y','Constant',true,'R2','Y');
Ecoli = Ecoli.add_regulator('Activation_out','R2','Y','I2');
Ecoli = Ecoli.add_regulator('Activation_in','G','U');
Ecoli.set('U','Constant',true,'G','U')
Ecoli = Ecoli.add_regulator('Activation_out','G','U','I1');
```

#### 2.4 CRISPR model

To achieve comparable concentrations between the Santos-Moreno model [3] and the Elowitz model, we fine-tuned several parameters, as detailed in Table S4 of the manuscript. Since in the Santos-Moreno model, the dilution rate $k_d$ is accounted for separately, we modified the original reaction constants for the degradation of proteins and RNAs by incorporating this parameter: $d_P = 0.0693 - k_d$ and $d_{RNA} = 0.3466 - k_d$. Additionally, the rate law for $R_1$ was slightly adjusted from the original Elowitz model to explicitly consider the DNA concentration ($[\mathrm{DNA}]_0 = 30$ molecules), resulting in the parameter correction: $a_1 = 30/30 = 1$ minute$^{-1}$.

RNA production was scaled by a factor of $r = 22.7$ using the Elowitz parameters for $R_1$–$R_3$, and we increased the production rates in $R_4$ and $R_6$ ($k_{fds}$, $k_{fdsd}$) with this factor as well. The Santos-Moreno model introduced additional algebraic equations to maintain overall concentrations of dCas and DNA, leading to differential-algebraic equations (DAE). In our modified model, we adjusted $R_5$ and $R_6$ to produce these species during dilution, thus ensuring mass conservation for DNA and dCas and converting the system to ordinary differential equations (ODE).

**Table S4:** Modified model for the CRISPRlator. Initial concentrations are set at $[\mathrm{DNA}]_0 = 30$ molecules and $[\mathrm{dCas}]_0 = 1434$ molecules [3]. All rate constants are provided in units of *minute* and *molecule*. The repression of the $j$th node on the $i$th node occurs through the formation of the $\mathrm{DNA}_j$-$\mathrm{sgRNA}_i$ complex, as described in the $\mathrm{R}_6$ reaction.

| Nr. | Reaction | Rate law | Rate constant | Ref. |
|---|---|---|---|---|
| $R_1$ | $\emptyset \longleftrightarrow \mathrm{mRNA}_i$ | $r_1 = a_0 + a_1\,[\mathrm{DNA}_i]$ | $a_0 = 0.03$ | [5] |
| | | $r_{1r} = (k_d + d_{\mathrm{RNA}})[\mathrm{mRNA}_i]$ | $a_1 = 1$ | |
| $R_2$ | $\emptyset \longleftrightarrow \mathrm{sgRNA}_i$ | $r_2 = a_0 + a_1\,[\mathrm{DNA}_i]$ | $d_{\mathrm{RNA}} = 0.3286$ | [5] |
| | | $r_{2r} = (k_d + d_{\mathrm{RNA}})[\mathrm{sgRNA}_i]$ | $k_d = 0.018$ | |
| $R_3$ | $\emptyset \longleftrightarrow \mathrm{P}_i$ | $r_3 = k_{\mathrm{P}}\,[\mathrm{mRNA}_i]$ | $k_{\mathrm{P}} = 6.9315$ | [5] |
| | | $r_{3r} = (k_d + d_{\mathrm{P}})[\mathrm{P}_i]$ | $d_{\mathrm{P}} = 0.0513$ | |
| $R_4$ | $\mathrm{dCas} + \mathrm{sgRNA}_i \longleftrightarrow \mathrm{dCas{:}sgRNA}_i$ | $r_4 = k_{\mathrm{fds}}[\mathrm{dCas}][\mathrm{sgRNA}_i]$ | $k_{\mathrm{fds}} = 1.4674$ | [3] |
| | | $r_{4r} = k_{\mathrm{rds}}[\mathrm{dCas{:}sgRNA}_i]$ | $k_{\mathrm{rds}} = 0.0776$ | |
| $R_5$ | $\mathrm{dCas{:}sgRNA}_i \longrightarrow \mathrm{dCas}$ | $r_5 = k_d[\mathrm{dCas{:}sgRNA}_i]$ | | [3] |
| $R_6$ | $\mathrm{dCas{:}sgRNA}_i + \mathrm{DNA}_j \longleftrightarrow \mathrm{dCas{:}sgRNA}_i{:}\mathrm{DNA}_j$ | $r_6 = k_{\mathrm{fdsd}}[\mathrm{dCas{:}sgRNA}_i][\mathrm{DNA}_j]$ | $k_{\mathrm{fdsd}} = 0.2670$ | [3] |
| | | $r_{6r} = k_{\mathrm{rdsd}}[\mathrm{dCas{:}sgRNA}_i{:}\mathrm{DNA}_j]$ | $k_{\mathrm{rdsd}} = 0$ | |
| $R_7$ | $\mathrm{dCas{:}sgRNA}_i{:}\mathrm{DNA}_j \longrightarrow \mathrm{dCas} + \mathrm{DNA}_j$ | $r_7 = k_d[\mathrm{dCas{:}sgRNA}_i{:}\mathrm{DNA}_j]$ | | [3] |

#### 2.5 The coherent and incoherent feed-forward loop

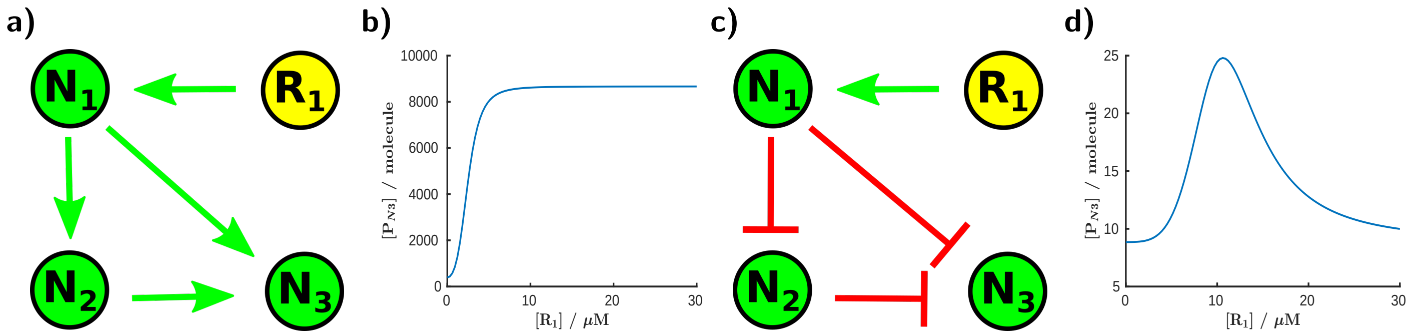


**Figure S6:** The behavior of coherent and incoherent feed-forward loops (CFFL and IFFL) with three nodes: $\mathrm{N}_1$, $\mathrm{N}_2$, and $\mathrm{N}_3$, where $\mathrm{R}_1$ serves as the inducer. The protein concentration produced by the third node, $\mathrm{P}_{\mathrm{N3}}$, is calculated as a function of the inducer concentration. The graphs display: (a) the circuit of a CFFL, (b) the output protein concentration as a function of the inducer concentration, (c) the circuit of an IFFL, and (d) the output protein concentration as a function of the inducer concentration. The detailed information about the models are available in the SI, specifically in the files "CFFL.html" and "IFFL.html".

**Listing 3:** Building the model for a coherent feed-forfard loop and an incoherent feed-forward loop:

```
%% coherent feed-forfard loop
Ecoli = Ecoli.add_node('N1','type1');
```

```
Ecoli = Ecoli.add_node('N2','type1');
Ecoli = Ecoli.add_node('N3','type1');
Ecoli = Ecoli.add_regulator('Activation_in','N2','HILL','P_N1');
Ecoli = Ecoli.add_regulator('Activation_in','N3','HILL','P_N2');
Ecoli = Ecoli.add_regulator('Activation_in','N3','HILL','P_N1');
Ecoli = Ecoli.add_regulator('Activation_out','N1','HILL','R1');

%% incoherent feed-forward loop
Ecoli = Ecoli.add_node('N1','type1');
Ecoli = Ecoli.add_node('N2','type1');
Ecoli = Ecoli.add_node('N3','type1');
Ecoli = Ecoli.add_regulator('Activation_out','N1','HILL','R1');
Ecoli = Ecoli.add_regulator('Repression_in','N2','HILL','P_N1');
Ecoli = Ecoli.add_regulator('Repression_in','N3','HILL','P_N1');
Ecoli = Ecoli.add_regulator('Repression_in','N3','HILL','P_N2');
```

#### 2.6 Novel oscillator circuits

##### 2.6.1 The "actolator" family

In the case of the repressilator, even a single-node circuit can oscillate, albeit through a different mechanism. The one-node oscillator is known as the Goodwin oscillator [7]. Oscillations can arise from high non-linearity in the system and inherent delays [8], or from a positive feedback loop with lower non-linearity [9]. Similarly, the actolator can exhibit oscillations with a two-node circuit, but achieving this requires a higher degree of non-linearity, typically a Hill exponent of at least 6, while generally used Hill exponents are around 2 (see the Elowitz-model in Table S2 and Figure S7).

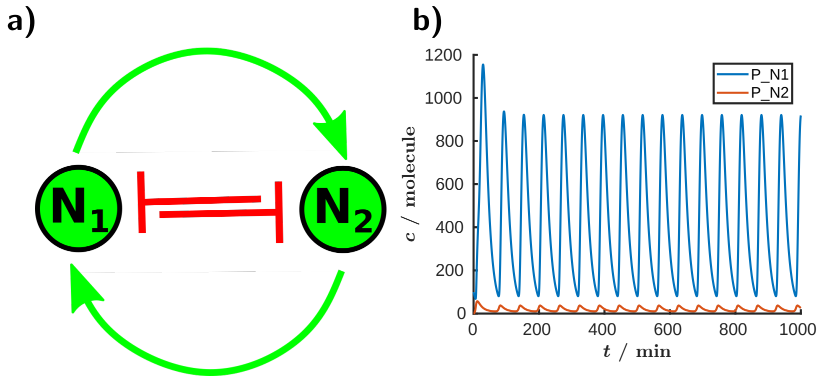


**Figure S7:** The "actolator" with a 2-node circuit. While the circuits in Figure 5 with 4, 6, 8, and 12 nodes exhibit oscillatory behavior with the original parameter set, the 2-node circuit required higher-order nonlinearity to induce oscillations. a) Circuit topology. b) Protein oscillations for the first (blue) and second node (red). Detailed information about the models is available in the SI, specifically in the file "actolator2.html".

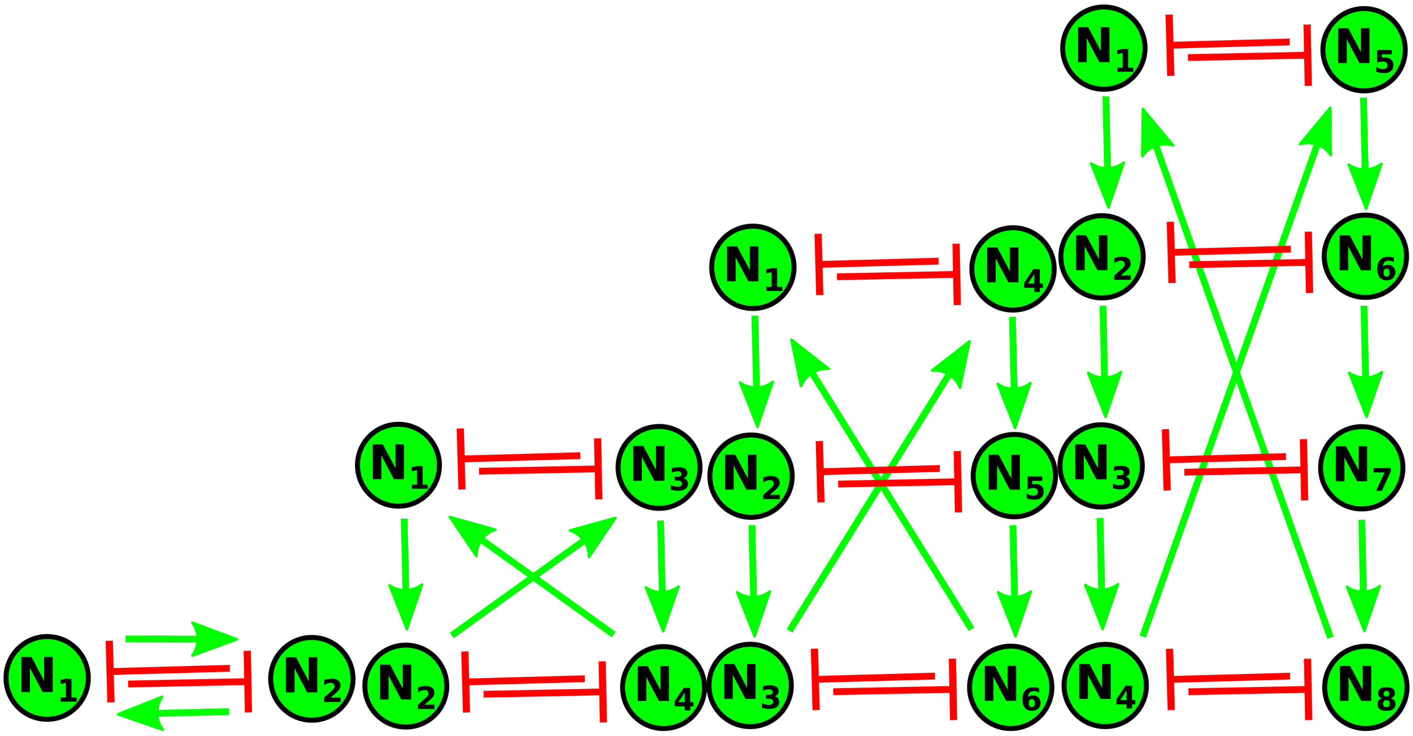


**Figure S8:** Alternative representation for the actolator family to highlight the role of the toggle switches. We show the 2, 4, 6, 8 node actolator respectively. In the case of the two node circuit, higher non-linearity is necessary for oscillation.

##### 2.6.2 Modified repressilator circuits: the "acrelator" family

In Figure S9 we demonstrate that when $n$ repressive interactions are replaced in an $N$-node circuit with activations, its oscillatory behavior resembles that of a repressilator with $N - n$ "effective nodes". If the number of "effective nodes" is odd, the circuit may exhibit oscillatory behavior. However, in many cases, achieving this requires a higher non-linearity, meaning a larger Hill exponent ($n_{Hill}$). For example, for a "x-y" circuit (Figure S11), $n_{Hill} \geq 6$ is needed, which is experimentally difficult to achieve, suggesting that such circuits may not oscillate in reality [10]. For a "4-1" circuit (Figure S10a), with a $n_{Hill} = 2$, the system exhibits damped oscillations, indicating that the fixed point is a stable focus. To achieve sustained oscillations, $n_{Hill} = 3$ was sufficient in this case, suggesting that this configuration might be experimentally feasible. We can observe a similar situation in the 6-node circuit: replacing one repressive interaction with activation enables oscillations with $n_{Hill} = 3$, while the "6-3" symmetrical circuit can oscillate even with the standard Hill exponent ($n_{Hill} = 2$). The "4-3" circuit, which includes one repression and three activation interactions, serves as a fundamental architecture for action potential oscillations. Indeed, this kind of circuit is not new to the oscillators field, and has been proposed to model the interactions between polarization, $Na^+$ flux, depolarization, and $K^+$ flux [10, 11].

It is worth discussing some additional interesting properties of these circuits, specifically the order of the nodes and their duty cycles. Due to the low half-saturation constants compared to the maximal protein concentrations, a node will activate almost simultaneously with its activating partner. Figure S11 illustrates this coordinated activation with the "4-3" circuit, where each node is activated nearly simultaneously. In a repressilator, the activation order of the nodes is straightforward: the next node to activate is the second one in the sequence, as two repressive interactions lead to one activation. When activations are also present in the circuit, they can be combined into a single node, causing these nodes to activate simultaneously. After this, the activation order can be determined using the same rules applied in the repressilator. However, we can observe notable differences in the duty cycles of the nodes. For example, in an activation chain like that in the "4-3" circuit, the first node in the chain will deactivate first, with subsequent nodes deactivating only afterward due to the low half-saturation constants. Consequently, the duty cycle will be longer for nodes positioned later in the chain, creating an asymmetry in the duty cycle that may be beneficial for certain applications. If a short duty cycle is desired during oscillations, selecting a node that is repressed by other nodes might be optimal. Conversely, for a node with a longer activation period, one connected through activation interactions would be preferable. Furthermore, nodes with a shorter duty cycle reduce the cellular burden.

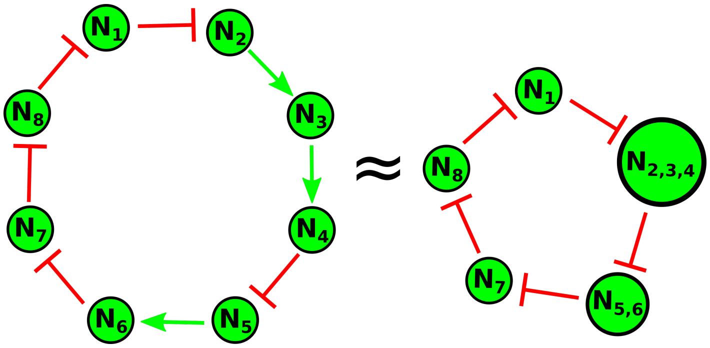


**Figure S9:** The "8-3 acrelator". Schematic representation of how oscillatory circuits with an even number of nodes can be created using only consecutive repressive or activating interactions, minimizing the number of regulatory steps. In this example, the eight-node circuit contains three activations, effectively reducing it to a five-node repressilator, which can exhibit oscillatory behavior. For simplicity, we refer to this circuit as the "8-3" circuit in the text. The different size of the nodes in the equivalent repressilator represents the asymmetry in the circuit, while multiple numbers in nodes – connected by activation – indicate that these nodes function synchronously. Further simulations of these types of circuits can be found in Figures S10 and S11.

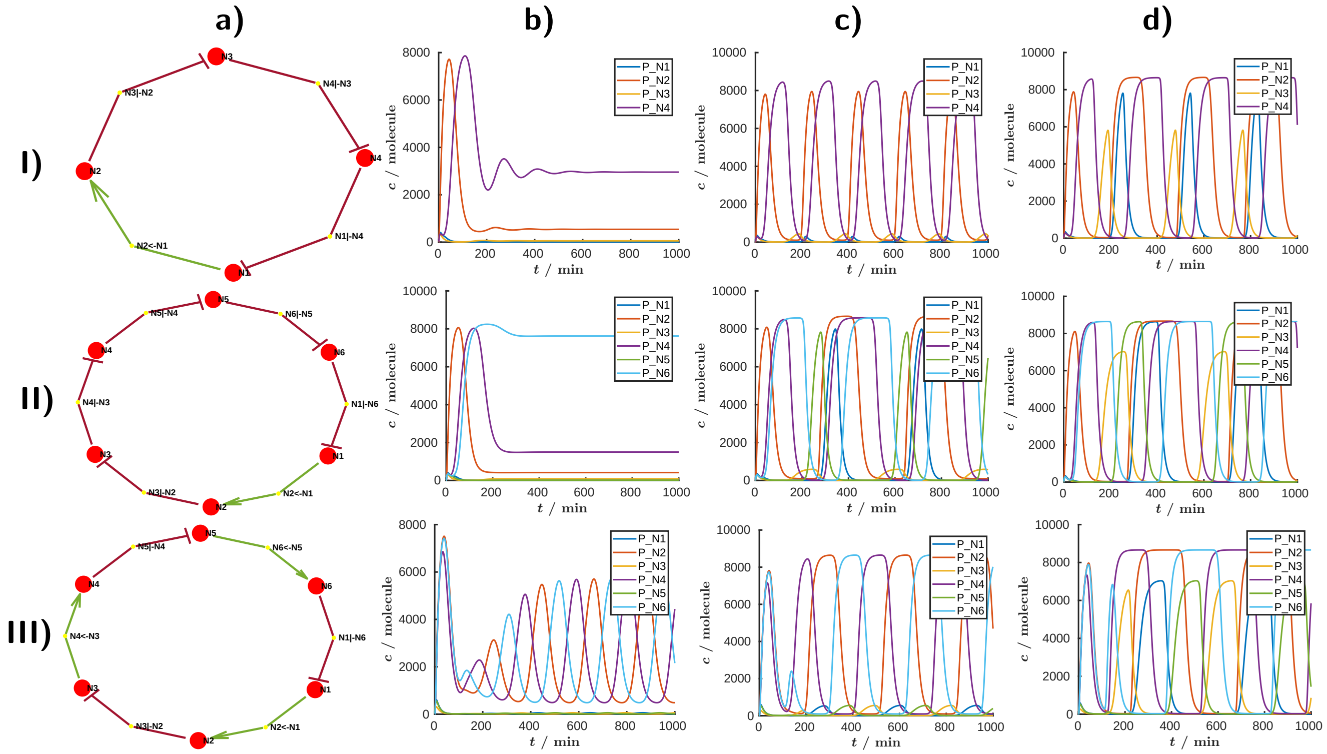


**Figure S10:** Examples of the "acrelator" family. These circuits are based on the repressilator family with an even number of nodes, where an odd number of edges have been replaced by activation interactions. I) A four-node circuit with one activation edge, II) a six-node circuit with one activation edge, and III) a six-node circuit with alternating repression and activation edges. a) Circuit topology, b), c), d) trajectories showing the protein concentrations over time for Hill exponents of 2, 3, and 4, respectively. Detailed model information is provided in the SI, specifically in the files "N4_A1.html", "N6_A1.html", and "N6_A135.html". Simulations were conducted using the repressilator model from [5].

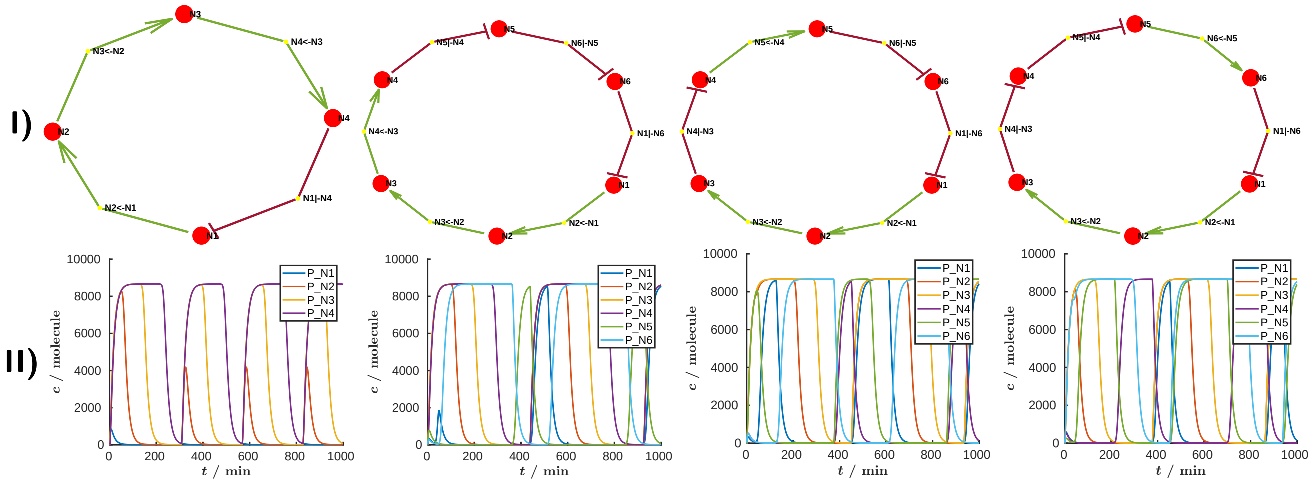

**Figure S11:** Examples of the "acrelator" family. These are based on the repressilator family with an even number of nodes, where an odd number of edges have been replaced by activation interactions. These circuits in this figure oscillate only with a higher Hill exponent, which was set to 6 for these simulations. I) Circuit topology, and II) the corresponding trajectories, showing protein concentrations over time. Detailed model information is available in the SI, specifically in the files "N4_A123.html", "N6_A123.html", "N6_A124.html", and "N6_A125.html". Simulations were performed using the repressilator model from [5].

#### 2.7 Light biosensor

**Table S5:** Fitted parameters for the light system in Eq. 1 to the experimental data shown Figure7a. The fitting was performed using MATLAB's `lsqnonlin` function, with a function tolerance of $10^{-14}$ and a termination tolerance of $10^{-10}$ for the independent variable. All parameters were fitted in logarithmic form, except for the Hill exponents.

| Parameter | Value | Unit | Parameter | Value | Unit |
|---|---|---|---|---|---|
| $k_{light}$ | 0.3185 | dimensionless | | | |
| $K_{light}$ | 156.9 | % | $K_{ara}$ | $2.756 \cdot 10^{-4}$ | % |
| $n_{light}$ | 0.9897 | dimensionless | $n_{ara}$ | 1.1177 | dimensionless |

To implement double-input regulations, new regulation types must be integrated into the node models. We illustrated this process using this example in Figure S12. Looking ahead, we also plan to develop a dedicated application to support the node model construction process.

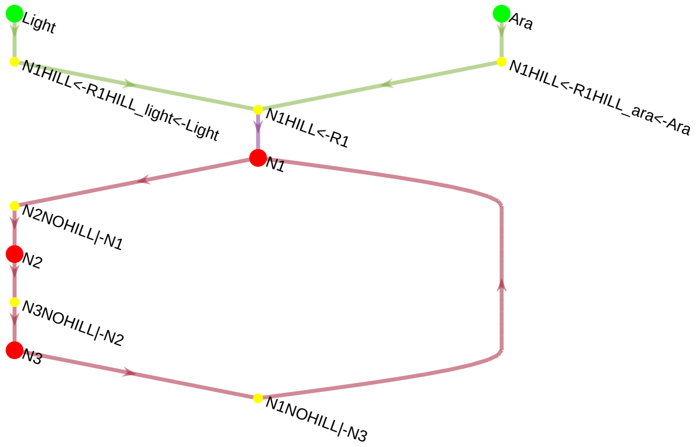


**Figure S12:** The three-node CRISPRlator circuit has two inputs: light and arabinose (Ara). The three nodes, N1, N2, and N3, repress each other consecutively through CRISPR interactions, with this input named "NOHILL". The N1 node is activated by N1HILL<-R1, which is activated by both light (N1HILL<-R1HILL_light<-Light) and arabinose (N1HILL<-R1HILL_ara<-Ara). This figure was created using the `make_graph()` method of the application (with a layered layout) and serves as an example of how to create multiple inputs for a node.
